# Deforestation projections imply range-wide population decline for critically endangered Bornean orangutan

**DOI:** 10.1101/2021.07.16.451448

**Authors:** Maria Voigt, Hjalmar S. Kühl, Marc Ancrenaz, David Gaveau, Erik Meijaard, Truly Santika, Julie Sherman, Serge A. Wich, Florian Wolf, Matthew J. Struebig, Henrique M. Pereira, Isabel M.D. Rosa

**Affiliations:** German Centre for Integrative Biodiversity Research (iDiv) Halle – Jena – Leipzig, Deutscher Platz 5e, 04103 Leipzig, Germany; Max Planck Institute for Evolutionary Anthropology, Deutscher Platz 6, 04103, Leipzig, Germany; Durrell Institute of Conservation and Ecology (DICE), School of Anthropology and Conservation, University of Kent, Canterbury CT2 7NR, UK; Borneo Futures, Bandar Seri Begawan, Brunei Darussalam; HUTAN-Kinabatangan Orang-utan Conservation Programme, Sandakan, Sabah, Malaysia; Center for International Forestry Research, P.O. Box 0113 BOCBD, Bogor 16000, Indonesia; TheTreeMap, Bagadou Bas, 46600 Martel, France; The University of Queensland, School of Biological Sciences, Brisbane, QLD, Australia; Natural Resources Institute (NRI), Agriculture, Health and Environment Department, University of Greenwich, Chatham Maritime, UK; Wildlife Impact, PO Box 31062, Portland, OR 97231, USA; School of Biological and Environmental Sciences, Liverpool John Moores University, Byrom Street, Liverpool, L3 3AF, United Kingdom; Institute for Biodiversity and Ecosystem Dynamics, University of Amsterdam, Science Park 904, 1098 XH Amsterdam, The Netherlands; CIBIO/InBio, Centro de Investigação em Biodiversidade e Recursos Genéticos, Laboratório Associado Universidade do Porto, Vairão, Portugal; CEABN/InBio, Centro de Ecologia Aplicada “Professor Baeta Neves”, Instituto Superior de Agronomia Universidade de Lisboa, Lisbon, Portugal; School of Natural Sciences, Bangor University, Bangor, Gwynedd, LL57 2DG, UK

**Keywords:** Biodiversity hotspots, Density distribution model, Future forest loss, *Pongo pygmaeus*, Tropics, Southeast Asia

## Abstract

Assessing where wildlife populations are at risk from future habitat loss is particularly important for land-use planning and avoiding biodiversity declines. Combining projections of future deforestation with species density information provides an improved way to anticipate such declines. Using the endemic and critically endangered Bornean orangutan (*Pongo pygmaeus*) as a case study we applied a spatio-temporally explicit deforestation model to forest loss data from 2001-2017 and projected future impacts on orangutans to the 2030s. Our projections point to continued deforestation across the island, amounting to a loss of forest habitat for 26,200 (CI: 19,500–34,000) orangutans. Populations currently persisting in forests gazetted for industrial timber and oil palm concessions, or unprotected forests outside of concessions, were projected to experience the worst losses within the next 15 years, amounting to 15,400 (CI: 12,000–20,500) individuals. Lowland forests with high orangutan densities in West and Central Kalimantan were also projected to be at high risk from deforestation, irrespective of land-use. In contrast, most protected areas and logging concessions currently harboring orangutans will continue to face low levels of deforestation. Our business-as-usual projections indicate the importance of protected areas, efforts to prevent the conversion of logged forests for the survival of highly vulnerable wildlife, and protecting orangutan habitat in plantation landscapes. The modeling framework could be expanded to other species with available density or occurrence data. Our findings highlight that species conservation should not only attempt to act on the current situation, but also be adapt to changes in drivers to be effective.

## INTRODUCTION

Borneo is globally important for biodiversity but experiences some of the highest deforestation rates in the world (Gaveau et al., 2016). As a consequence of agriculture, mining, infrastructural development and forest fires, the area of old-growth forest on Borneo declined by 14% between 2000 and 2017 (Gaveau et al., 2018). To allow for development while reducing deforestation pressures on the natural environment, land-use planning and conservation should better incorporate insights from past patterns and drivers of land-use change, and consider how deforestation could continue into the future.

Advances in spatially-explicit and dynamic deforestation modeling offer new ways to study current and expected future forest loss in the tropics (Rosa et al., 2013). In comparison to previous approaches, these models dynamically project deforestation as a sum of local events, influenced by past patterns of various drivers, rather than imposing a fixed deforestation rate based on historical trends. While deforestation projections have been more commonly applied in the South/Central tropical region (Rosa et al., 2013; Silva et al., 2020), there are far fewer assessments available for Southeast Asia despite this being a region of high forest loss.

Recent increases in the availability of species observation data, as well as advances in computational power and statistical methods, provide improved estimates of range-wide species density distributions (e.g., Strindberg et al., 2018; Wich et al., 2016). A density distribution model for the Bornean orangutan (*Pongo pygmaeus*), for example, indicated that the population declined by >100,000 individuals between 1999 and 2015 (Voigt et al., 2018). Large-scale deforestation, together with killing in conflict or for food, severely threatens the long-term population viability of this species and stable orangutan populations only persist in landscapes with sufficient forest cover (Ancrenaz et al., 2016).

Here we use the Bornean orangutan as a case-study to demonstrate how coupling of deforestation projections with density distribution models can help estimate future population impacts of land-cover change on Critically Endangered and forest dependent species. We tailored a deforestation model to each Bornean administration within the orangutan range (five Indonesian provinces; two Malaysian states - hereafter all referred to as provinces), identified drivers and patterns of land-cover change in the past (2000-2017) and projected them into the future (2018-2032) under a business-as-usual scenario. By identifying the population units most vulnerable to potential future deforestation our approach can be used to guide pre-emptive conservation efforts and serve as a business-as-usual baseline against which certain policy interventions could be tested. The approach could be equally as valid for other species and regions where wildlife information and deforestation trends are well documented.

## METHODS

### Forest maps and deforestation drivers

We utilized mapped forest trend data specific to Borneo, which quantifies forest loss between 2001 and 2017 at a resolution of 30 m (CIFOR (2019); Table 1). Forest loss or deforestation is defined as the annual removal of intact or logged old-growth forest that is closed-canopy, high-carbon evergreen dipterocarps on mineral or peat soils (>90% cover), low-biomass pole forests on peat domes, heath forest, and mangroves (CIFOR 2019). This definition matches the “primary” and “secondary” forest definition of the Indonesian government (Ministry of Environment and Forestry Republic of Indonesia, 2018). The binary forest and forest loss layers were resampled to a 1 km^2^ pixel size using nearest-neighbor resampling.

**Table 1:** Predictors used in deforestation modelling, including their description, source and year (Processing information in Supporting Information S1).

| Name | Description | Source | Year |
| --- | --- | --- | --- |
| Forest loss | Forest loss previous to calibration period (2001-2012) and in calibration period (2013-2017)<sup>a</sup> | Centre for international Forestry Research (2019), Gaveau et al. (2018) | 2001-2012, 2013-2017 |
| Elevation | Elevation in meters derived from digital elevation model | Jarvis et al. (2008) | 2000 |
| Distance to roads | Distance to primary and logging roads | Center for International Earth Science Information Network - CIESIN - Columbia University (2013), Gaveau et al. (2014) | 2010, 2013 |
| Distance to rivers | Distance to major rivers with a minimum of 200 km<sup>2</sup> drainage area | Abram et al. (2015) | 2010 |
| Active fire incidence | Aggregated number of active fires (MODIS and VIIRS) | MODIS Collection 6 NRT (2018), VIIRS 375m NRT (2018) | 2000/2002-2017 |
| Human population density | Number of humans within 1 km<sup>2</sup> | Bright et al. (2012) | 2012 |
| Land-use and management | Including protected areas, logging concessions, industrial timber plantation concessions, industrial oil palm plantation concessions, unprotected areas outside concessions (as reference areas)<sup>b</sup> | IUCN & UNEP-WCMC (2017) Santika et al. (2015) | 2012, 2017 |

<sup>a</sup> Forest loss previous to calibration period was used to inform the projection for the calibration period, while forest loss in the calibration period was used to inform projections in the future. <sup>b</sup> Areas outside of protected areas or concessions were included in land-use and management as a reference area, i.e., it was coded as level 0.

Patterns of tropical deforestation are shaped by physical and accessibility characteristics, anthropogenic pressures, and land-use (Austin et al., 2019; Curtis et al., 2018). We compiled data on elevation, distance to roads and rivers, human population density, occurrence of fire, and land-use and management as spatial predictors (Table 1). The selection was based on literature describing important drivers of deforestation in the tropics and for Borneo specifically (Austin et al., 2019; Rosa et al., 2013; Struebig et al., 2015). While forest was aggregated for the calibration period (2013-2017) and the years preceding the interval (historic deforestation: 2001-2012), fire occurrence was aggregated over the available time. For the remaining predictors the available time point closest to the calibration interval was chosen (Table 1).

Borneo is governed by multiple countries, each with their own land-use system. We amalgamated these systems into the following land-use types: protected areas, logging concessions, industrial timber and oil palm plantations, and areas not allocated to protection or concessions (i.e. areas without any formal management, as well as urban or infrastructure development areas) (Santika et al., 2015). Protected areas were based on the WDPA database (IUCN and UNEP-WCMC, 2017) and national land-use plans (as used in Santika et al., 2015), and were further differentiated into three categories: areas listed in the WDPA database as IUCN category 1-3 were categorized as ‘strictly protected areas’; those listed as IUCN category 3-6 were categorized as ‘sustainable use protected areas’; and the remainder as ‘national protected areas’. (Table 1; Supporting Information S1). Land-use classes describe the designation and not the land-cover, hence concessions can include forests that have not yet been converted or logged.

All layers were converted to the Asia South Albers Equal Area Conic projection and resampled to the same extent and origin at 1 km^2^ pixel size, the highest resolution common to all layers, using bilinear interpolation for continuous predictors and nearest-neighbor interpolation for categorical predictors. All spatial manipulations were performed in Python (Python, 2016), and aggregated, analyzed and visualized in Python, R (R Core Team, 2020) and ArcGIS (Esri Inc., 2014) (Supporting Information S1 for full processing details).

### Deforestation model framework

We used the modeling approach developed by Rosa et al. (2013) for each Bornean province to project the probability of deforestation into the future. The model accounts for stochasticity of deforestation events, and province-wide forest loss rates emerge as the sum of local scale deforestation events, resulting from the influence of drivers operating in each particular province. The model is based on the probability that trees in a pixel are lost in a certain time interval. Using a forward stepwise regression, models were fitted to five years of forest loss data from 2013–2017 (calibration period). We selected this calibration interval length by considering the trade-off between short intervals, potentially reflecting exceptional years, or long intervals, potentially including outdated trends, as recommended by Rosa et al. (2015).

To assess the predictive power gained by adding predictors to the model, a cross-validation technique was used. This technique allowed us to check how accurately the model projected deforestation with the added predictor compared to a randomly selected subset of 50% of the data that was not used to train the model. After successively adding the variable that resulted in the highest likelihood model, the overall best model out of a total of 31 models was selected for each province individually (Table S2, Supporting Information S2).

### Simulations

After testing the predictor combinations for each province, the model with the highest test likelihood was used to project the probability of deforestation for each pixel in the five-year calibration period (2013–2017) and the following three five-year periods (2018–2022, 2023– 2027, 2028–2032).

The simulation was based on updating the model for each iteration and time step. Predictor uncertainty was incorporated by drawing the values for the simulations from a Gaussian distribution, using the estimated mean and standard deviation. We subsequently evaluated whether or not a pixel in a certain period and for a certain iteration was lost, by comparing its probability of deforestation with a randomly drawn number from a uniform distribution between 0 and 1. We then classified the pixel as deforested if the number was less than the probability of deforestation. This procedure was repeated for all four time steps and run multiple times (n = 100) to gauge uncertainty in model predictions. The generated binary forest maps were used to calculate projected deforestation and impact on orangutan populations. Furthermore, to characterise the deforestation risk across provinces and land-use classes, the binary maps were aggregated into a summed probability of deforestation. This value represents the fraction of simulation runs in which the forest in a pixel was lost; i.e. if a pixel was selected to be deforestation in that time period in 50 out of 100 iterations, then it has a 50% probability of deforestation.

### Validation and analysis

We validated the models for each province against observed data for the calibration time-period (2013–2017), by calculating the area under the Receiver Operating Characteristic (ROC) curve (AUC value) for the 100 iterations. We also calculated the proportion of match between observed and cumulative forest loss within certain distances (0, 1, 5 and 10 km) surrounding the pixel following Rosa et al. (2013) for each province and the whole island.

### Impacts of projected deforestation on orangutan abundance

We calculated the projected future impact of deforestation on orangutans by overlaying the projected forest loss and summed probability of forest loss with current orangutan density distribution maps. Orangutan density distribution was based on orangutan nest surveys implemented between 1999 and 2015 (4,316 km survey effort, a median of 86 transects per year) and a predictive density distribution model. The model considered survey year, climate, habitat cover and human threat predictors to estimate range-wide patterns of orangutan abundance and is described in full in Voigt et al. (2018). We generated a baseline orangutan distribution for 2018 by excluding pixels deforested until 2017 from the density distribution layer of 2015.

To estimate the total projected loss of orangutans as a consequence of deforestation we excluded all pixels with forest loss from the baseline orangutan abundance map based on the binary maps of projected deforestation, and summed the number of affected orangutans. Vulnerability of orangutan populations was assessed by calculating the proportion of orangutan numbers within pixels with either low (0–33%), medium (≥ 33–67%) or high (≥ 67–100%) summed forest loss probability. Orangutan abundance was also classified into low (0.01–0.5 individuals/km^2^), medium (>0.5–2 individuals/km^2^) or high (> 2 individuals/km^2^) local orangutan abundance. Abundance thresholds were based on the spread of local densities and expert assessment of what constitutes low, medium or high orangutan density throughout Borneo (Utami-Atmoko et al., 2019). Last, we calculated the loss of forest and vulnerability and loss of orangutans within provinces and land-use categories. Confidence intervals of the number of orangutans affected were generated by randomly pairing deforestation projections (n=100) with bootstraps of orangutan abundance (n=1000) (Voigt et al., 2018). All orangutan numbers were rounded to the nearest 100.

## RESULTS

### Deforestation model for Borneo

In all provinces, previous forest loss, distance to roads and land-use were included in the best model (Table S1). Distance to rivers and elevation were included for six of the seven provinces, fire incidence for five provinces, and population density for three provinces. Probability of deforestation was highest near areas of past forest loss (Figures 1 and 2e).

**Figure 1:**
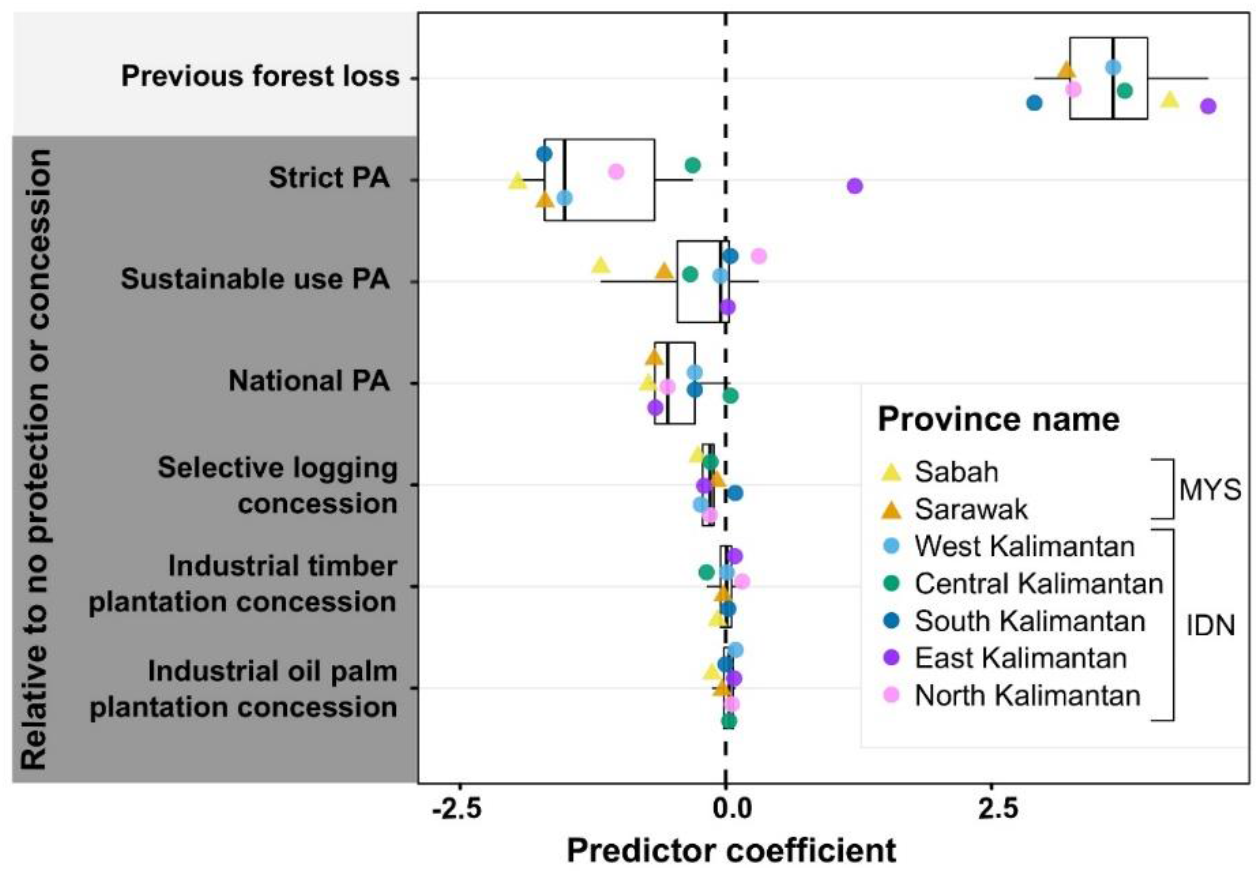
Influence of land-use and management predictors across Malaysian and Indonesian provinces on Borneo. Model coefficient values across provinces are summarized in a boxplot (median and 25^th^ and 75^th^ quartiles as hinges). Predictors with a coefficient smaller than zero (dashed line) were related to lower forest loss, while predictors with a coefficient larger than zero to higher forest loss. The effect of protected areas (PA) and concessions (grey shaded background) is relative to the effect of no protection or designation as concession. Strict PAs are IUCN category 1-3, sustainable use PAs are IUCN category 3-6 or no category and all protected areas recognized in the national land-use plans but not represented in the WDPA database (2017) are included as national PAs (Supporting Information S1 and S2). The intercept and predictors for which all provincial coefficients were close to zero (mean absolute coefficient smaller than 0.05 and a spread smaller than 0.1) were excluded from the figure (elevation, distance to road and rivers, fire incidence, human population pressure). The 95% confidence intervals derived from the 100 model iterations around points are not shown, as they fall within the points. IDN–Indonesia, MYS–Malaysia.

**Figure 2:**
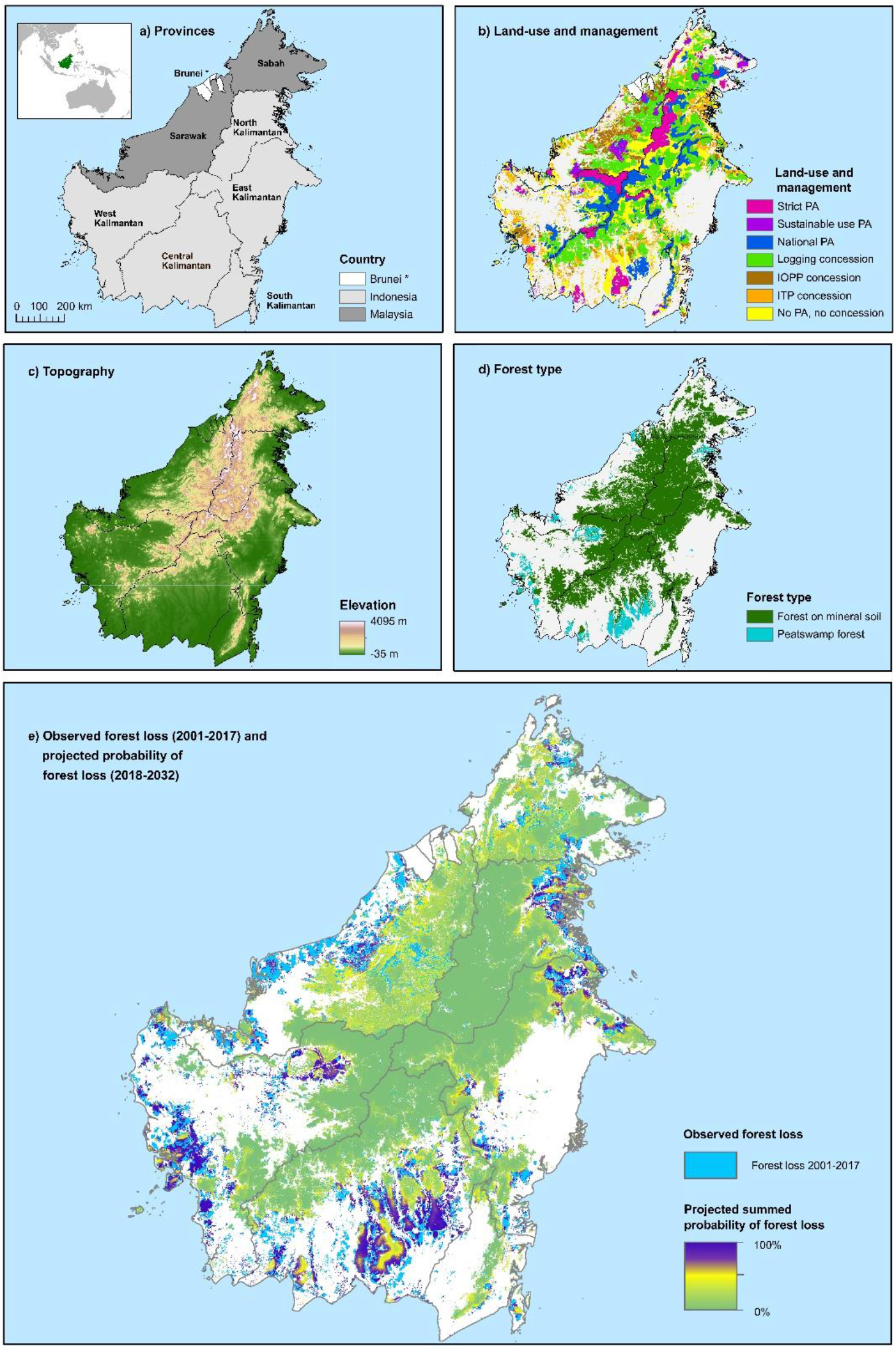
Projected deforestation probability and contextual layers across Borneo. a) Administrative boundaries of Indonesia, Malaysia and Brunei. The position of Borneo can be seen in the inlay. Brunei is excluded from maps b and e, as important predictors did not contain sufficient information for this country. b) Land-use and management within forested areas (PAs–Protected areas, ITP–industrial timber plantation, IOPP–Industrial oil palm plantation). c) Elevation was derived from a digital elevation model by Jarvis et al. (2008), d) Forest types were derived from Miettinen et al (2016) by combining lowland, lower montane and upper montane evergreen forests to represent forests on mineral soils. g) Observed deforestation and projected probability of forest loss on Borneo over time (2018–2032). Observed deforestation and the individual projection time steps are shown in Figure S2.

Protected areas experienced low levels of projected deforestation, with the lowest levels associated with strictly protected areas (Figure 1). Logging concessions were associated with lower probability of forest loss, with the exception of concessions in South Kalimantan. Industrial timber and oil palm plantation concessions had similar levels of deforestation compared to areas without formal management. Although included in the best models of provinces, elevation, distance to roads or rivers, fire incidence and population density were weak deforestation predictors, with model effect sizes close to zero.

### Model validation

The model selection process yielded provincial models with good discriminatory power with mean AUC values ranging from 0.80 (Sarawak) to 0.92 (North Kalimantan) (standard deviation: 0.0003–0.0016). A median of 17.3% (Interquartile range (IQR): 9.58%) of all pixels projected to be deforested were in the exact locations of observed forest loss in the calibration period (Figure S1). However, 56.3% (IQR: 15.3%) of pixels were in the direct neighborhood (within 1 km) and 95.8% within 5 km (IQR: 2.56%) of observed deforestation, indicating strong spatial match between projections and observed values.

### Spatio-temporal deforestation and projections

Between 2000 and 2017 forests on Borneo decreased by 59,949 km^2^, and by 2032 a further 74,419 km^2^ (95% confidence interval (CI) 74,023–75,157 km^2^) was projected to be lost - a 32% decrease since 2000 (Figure 2 and 3, Table S2 and Figure S3). Past annual deforestation rates, measured in percent forest lost relative to forest cover in 2000, ranged between 0-3% for all provinces, with high inter-annual fluctuations (Figure 3b). Projected median annual deforestation rates (2018-2032) ranged between 0.55 and 1.72% (Figure 2).

**Figure 3:**
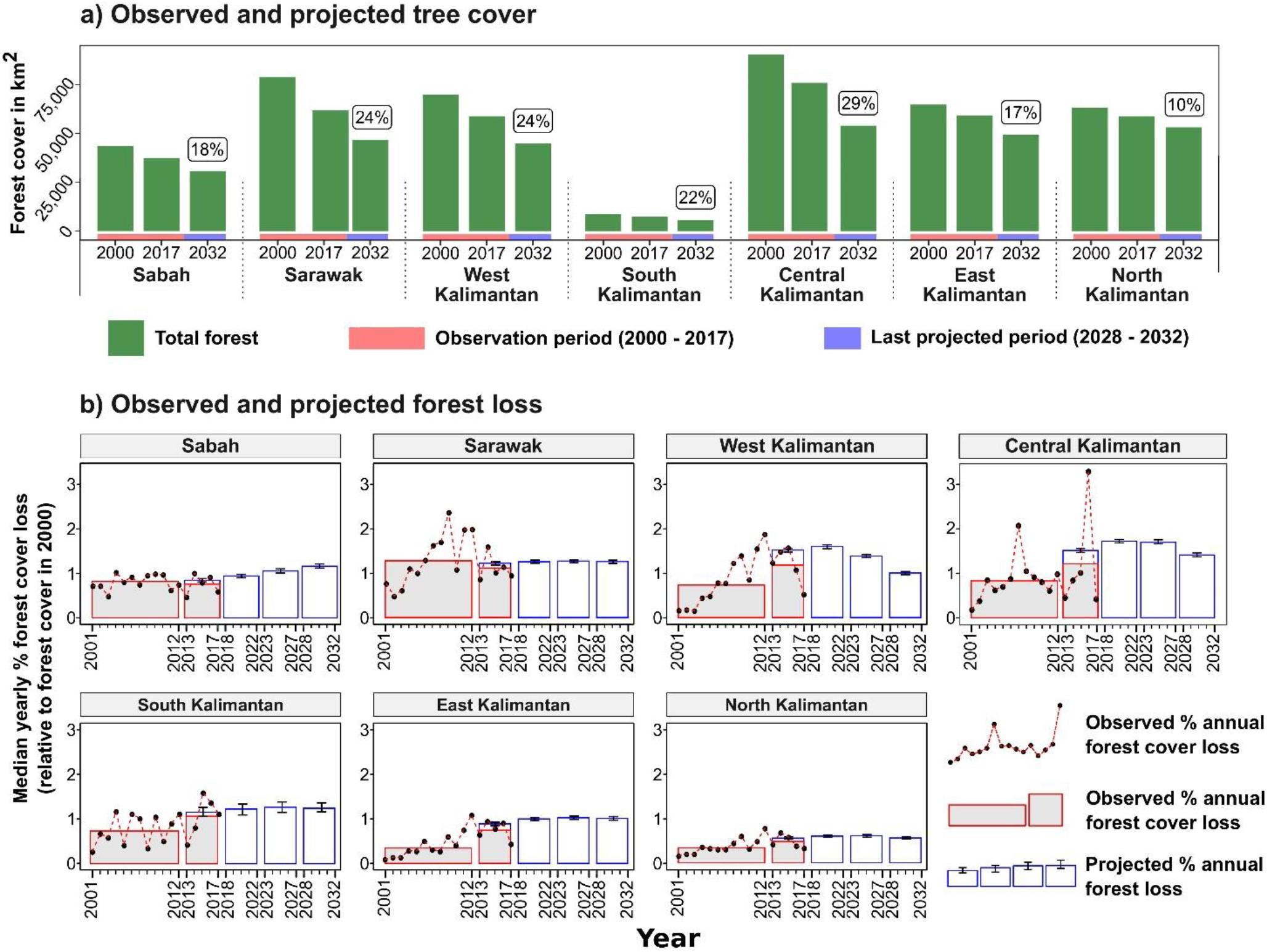
Observed and projected forest area and loss across Borneo from 2000 to 2032 a) The total forest in the first and last year of the observation period (2000 – 2017, red axis) and the median forest in the last projected five-year period (2028 – 2032, blue axis) for each province (95% confidence interval (CI) as error bars). Percent future forest loss from 2018 to 2032 is given above the bars (CI in Table S3). b) Aggregated average percent forest loss before simulation (2001 – 2012) and in the calibration period (2013 – 2017) (red bars with grey filling) was used for model fitting. The annual observed forest loss (red line with black dots) shows inter-annual variability of forest loss in the provinces. Deforestation was simulated for the calibration period and three five-year periods from 2013 – 2032 (blue bars, n = 100, error bars represent CI). The calibration period from 2013-2017 can be compared to the projection of forest loss in the same time interval (difference presented in Table S3). All values in b) given in annual percent loss of forest in 2000, by aggregating over the time-period over which the bar extends and dividing by number of years in interval.

At the provincial level, projected loss of forest area ranged from 10% in North Kalimantan to 29% in Central Kalimantan in comparison to forest in 2017 (Figures 3b and Table S2). Deforestation trends tended to vary among provinces because of differences in drivers and their relationship with deforestation, as well as the distribution of clusters with high deforestation probabilities (Figure 2e and Figure 3a). The deforestation rate was projected to increase over time and then stabilize in Sarawak, South and East Kalimantan, decrease in (2028-2032) in West Kalimantan, Central and North Kalimantan, and continue to increase at a low level in Sabah (Figure 3b). In the calibration interval the projected median annual loss was larger than the observed rate, with a deviation between 0.07% (North Kalimantan) to 0.33% (West Kalimantan) (Table S3). However, in all provinces the projected median deforestation rate was within the range of the observed annual rates, indicating a good fit of projections.

Across provinces, protected and high-elevation areas had a high probability of maintaining forest cover until 2032 (Figure 2). Lowland forests, those within industrial timber and oil palm plantations, and forests without protection or concession status, were all associated with a low probability of maintaining forest cover and a high vulnerability to future deforestation.

### Orangutan vulnerability in provinces

Medium to high (> 0.5 ind/km^2^) orangutan abundances are concentrated in the protected lowlands and peatswamp forests in West, Central and East Kalimantan as well as the forests at higher elevations along the border of West and Central Kalimantan. High local orangutan abundances (> 2 ind/km^2^) coincide with high risk of deforestation (i.e. summed probability of projected deforestation ≥ 67%) in the unprotected lowland and peatswamp forests of West, Central and East Kalimantan. In contrast, areas with medium to high orangutan abundance in the central part of West and Central Kalimantan at higher elevations had low deforestation probability (< 33% summed probability of projected forest loss) (Figure 4a).

**Figure 4:**
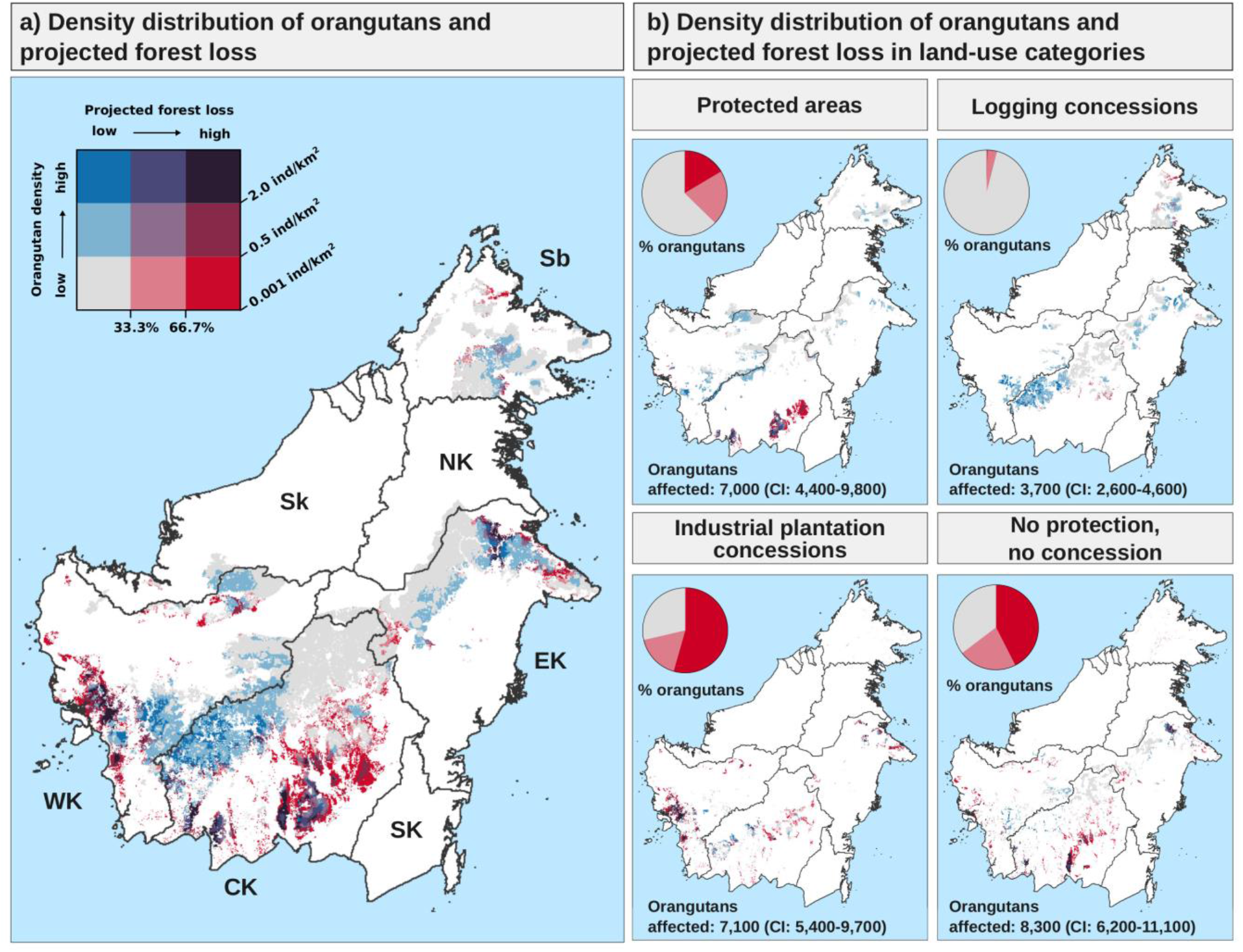
Density distribution of orangutans and summed probability of projected forest loss in land-use and management areas until 2032. The density of orangutans is indicated by blue shades and the probability of forest loss by red shades (individual maps in Figure S3). Darker colors identify higher levels of orangutan density and summed probability of projected deforestation. b) Forest of strict, sustainable use, and national protected areas were aggregated to a single category. Similarly, industrial timber and oil palm plantations concessions were combined into a single industrial plantation concession class. The proportion of orangutans in areas with low, medium or high levels of forest loss (pie charts, red shades only) and total projected loss of orangutans until 2032 (number in each panel) differed between land-use classes. Numbers shown are rounded to the nearest 100. Only pixels that were forested in 2017 and that have an estimated density of >0.001 orangutans/km^2^ are represented.

Although fewer orangutans occur in Sabah and Sarawak compared to other provinces, most are projected to experience low levels of forest loss (Figure 4 and Figure S4). In these two states only 9% (Sabah) and <1% (Sarawak) of orangutans occurred in areas with high deforestation probabilities. Conversely, in West, Central and East Kalimantan 27%, 23% and 15% of all orangutans were in areas with high deforestation probabilities (Figure S3). Orangutans are only present in very low numbers or entirely absent from North and South Kalimantan.

### Orangutan vulnerability in land-use and management categories

Orangutans within protected areas and logging concessions were found to be less vulnerable to deforestation than orangutans in industrial plantations and in areas without management. Overall, forests in protected areas and logging concessions harbored 68% (CI: 65–70%) of all orangutans estimated to occur on Borneo in 2018. Most of these orangutans inhabited forests with low deforestation probabilities: 62% (CI: 52–72%) of all orangutans within protected areas and 96% (CI: 95–97%) within logging concessions. Nevertheless, deforestation was projected to affect 7,000 (CI: 4,400–9,800) orangutans in protected areas and 3,700 (CI: 2,600–4,600) orangutans in logging concessions, amounting to 27% [CI: 22-31%] and 14% [CI:11-17%] of all orangutans lost, respectively.

Conversely, a large percentage of the orangutans inhabiting forests allocated for industrial plantations depended on habitat that was highly susceptible to deforestation. Combined these could affect 7,100 orangutans (CI: 5,400–9,700), representing 27% (CI: 25-31%) of the loss of orangutans on Borneo.

Areas without formal management supported 19% (CI: 18–21%) of all orangutans in Borneo, and much of this was at high risk of deforestation according to projections (23% [CI: 22–26%] medium, 44% [CI: 37–51%] high risk areas), affecting 8,300 (CI: 6,200–11,100) orangutans (32% [CI:31—32%)] of all loss). Those areas with high vulnerability also harbored high orangutan densities, notably around the vast Sabangau peatlands in Central Kalimantan and in the Lesan-Wehea landscape in East Kalimantan (Figure S5).

## DISCUSSION

Wildlife population management is informed by our knowledge about drivers of declines and our ability to anticipate which measures could effectively curb those losses. Orangutans, like many other tropical and forest dependent species, have been affected by deforestation in the past, and populations have declined dramatically (Ancrenaz et al., 2016). Our modelling of deforestation trends revealed that the forests of Borneo are projected to decline by a further 19% until 2032. Annual deforestation was projected to occur at a rate of 1.54%, which is similar to that experienced in Sumatra since 2001 (Gaveau et al. (2021a)), but higher than that reported from West Papua (0.82% between 2019 −2036, Gaveau et al. (2021b)) or Wallacea (1.23% between 2019, Voigt et al. (in review)). Rates across much of Indonesia remain higher than the pan-tropical average (0.49%) during the 1990s and 2000s (Achard et al., 2014).

Protected areas and logging concessions are associated with the lowest deforestation risk, in line with previous research on Borneo (Gaveau et al., 2018, 2013; Santika et al., 2015). The sizeable orangutan populations remaining in these areas are thus largely spared from deforestation. Rather, the greatest deforestation threats to orangutans remain in forests allocated for conversion to industrial timber and oil palm plantations, or those with no formal land-use designation. In these forests around 81% (CI: 78–85%) of the orangutan inhabitants could be lost, particularly in the peatlands of West and Central Kalimantan.

### Importance of protected areas and logging concessions

Although protected forests experienced lower deforestation overall and thus are effective in protecting orangutan habitat, some were projected to experience elevated deforestation in the future - most notably Sabangau national park, which currently contains high numbers of orangutans (Voigt et al., 2018). This region is particular vulnerable to peatland fires that have caused considerable forest loss in the past (Drake, 2015). Our future projections thus point to the relevance of continued measures and policies to prevent uncontrolled fires in general, and specifically in this highly important conservation area. Potential measures used in Indonesia include strengthening fire-management policies and fire-fighting efforts or enforcing fire bans (Carmenta et al., 2017). Restoring degraded peatland and providing incentive schemes for smallholders to comply with environmental legislation and manage their land without fire are key to minimizing further fire-induced deforestation in Kalimantan, as these habitats are particularly prone to burning (Santika et al., 2020).

Logging concessions harbored the largest proportion of Bornean orangutans in areas with low deforestation risk, and are thus expected to be important refuges for Bornean orangutan populations in the future. This is not surprising as logged forests on Borneo, although disturbed, tend to support comparable species numbers to unlogged forest (Deere et al., 2017), while containing the largest number of orangutans in the past (Voigt et al., 2018). Our findings reinforce the value of well-managed logging concessions for biodiversity and the need to control habitat degradation within these forests. Nevertheless, logged over forests throughout Borneo are at risk of degazettement and subsequent clearing due to decreased profitability after multiple logging cycles and long periods of inactivity. This makes concessions vulnerable to encroachment by smallholder and industrial agriculture, accelerating loss of forest in these areas (Burivalova et al., 2020).

### Disproportionate biodiversity losses in plantations and other non-protected areas

The largest conservation gains can be made by effectively curbing deforestation in and around plantations landscapes and non-protected forest areas. In forests allocated to industrial timber or oil palm, a large proportion of orangutan habitat is at elevated risk of deforestation. In particular, conversion of peatswamp forest to plantations in West and Central Kalimantan could lead to the loss of high orangutan numbers since these areas still support high orangutan densities. However, in Indonesia, the conversion of primary and peatswamp forests to plantations has been banned through a moratorium since 2011 (Widodo, 2017), and Sabah has also committed to protecting remaining forests (Sabah Forestry Department, 2017). Although the Indonesian moratorium excludes secondary forests impacted by selective timber harvest, the moratorium indeed seems to have slowed down deforestation rates in non-fire years (Chen et al., 2019).

Other tools available to slow deforestation in areas slated for conversion include corporate zero-deforestation pledges, which are gaining traction in the oil palm sector (https://rspo.org/news-and-events/news/uniting-to-deliver-deforestationfree-sustainable-palm-oil-more-critical-than-ever). Forest patches retained in plantation landscapes under such practices can provide valuable habitat for wildlife, including orangutans (Deere et al., 2020), although an impact evaluation found only moderate effects on avoided deforestation in Sumatra and Kalimantan prior to 2018 (Carlson et al., 2018). The greatest gains from zero deforestation pledges will come from companies not clearing any new forest areas in the first place, and the impact of this will take time to be detected. The implementation of such tools are thus useful to avoid loss of valuable orangutan habitat and maintain connectivity of forest areas within plantations, mitigating the projected impacts on orangutans in the future (Meijaard et al., 2017).

Our analysis also highlights the importance of areas that are not protected or allocated to concessions. Here, the largest number of orangutans (8,300, CI: 6,200–11,100, i.e., 32% [CI: 31–33%] of total projected decline) is estimated to be extirpated until 2032. By addressing deforestation drivers in these areas, considerable losses to orangutan populations could be prevented. However, the de-facto management is very heterogeneous in unallocated areas and individual conservation solutions need to be tailored to the local conditions to effectively alleviate deforestation risk.

### Modeling uncertainties, caveats and future development

With the presented deforestation projections, we created a business-as-usual baseline against which future developments in the Bornean orangutan range can be compared. Although it is likely that the deforestation in coming years has similar drivers and patterns than the deforestation in the recent past, it cannot be assumed that future dynamics will perfectly mirror the past. The effects of changes in political agendas and development priorities (e.g., Ferrante and Fearnside, 2019), fluctuation of commodity prices for important agricultural products (Gaveau et al., 2018), global climate changes and repercussion of events such as the global COVID-19 pandemic on forests (Brancalion et al., 2020) are difficult to anticipate and thus directly include in models.

To manage for this uncertainty, scenarios could explore potential future developments, such as investment in mines, dams and infrastructure projects, further agricultural expansion and the implementation and effectiveness of deforestation mitigation measures and their effect on orangutans.

However, as of yet the data to parameterize scenarios at the scale of Borneo are not freely available. Such data could be more easily compiled at the local scale, where relevant stakeholders could more readily co-develop realistic development scenarios relevant for their landscapes, and explore expected outcomes, including impacts on biodiversity. In this study we could show that drivers and patterns of deforestation vary for the different provinces, producing more regionally-relevant projected deforestation rates and highlighting the potential for models that are tailored to the local scale context to project future change.

This study omits a proportion of orangutans (ca. 1,800 individuals, 2.6% of all Bornean orangutans, unpublished analysis) potentially surviving in forest fragments within agricultural landscapes or other marginal habitat. Although orangutans depend on forest habitat, individuals have occasionally been found to persist in mosaics of forests and plantations (Ancrenaz et al., 2021; Seaman et al., 2019). Populations thus seem to persist in human-modified landscapes where there is connectivity with larger forest areas and no mortality. Crucially, however, our understanding of the habitat requirements that guarantee orangutan survival at densities sufficient to support populations, (e.g. minimum area requirements, food availability and landscape configuration), is not well established. Both the mapping of relevant features and knowledge of orangutan population dynamics within fragmented landscapes are in their infancy, and thus cannot yet be implemented in a modelling context to provide meaningful insights. These areas merit more attention within remote sensing and orangutan research agendas to improve species conservation in human-modified landscapes.

Orangutans are not only threatened by deforestation, but also suffer considerable declines through hunting, killing in conflict situations and live capture. These threats often remain hidden and are governed by complex socio-economic drivers that remain poorly understood and mapped, hindering rigorous spatial assessment (Meijaard et al., 2011). Although conflict killing and the capture of young orangutans as pets have been shown to increase close to recently converted forests (Santika et al., 2017), the threat remains relevant across converted and unconverted landscapes and impacts of hunting on Borneo are not well represented by accessibility or human population density. This makes it challenging to model the contribution of this threat to orangutan vulnerability (Meijaard et al., 2011; Sherman et al., 2020). The projected orangutan losses thus only represent a proportion of future population losses and are relatively conservative. Additional to measures implemented to curb deforestation, no-killing policies are an essential cornerstone of any conservation approach to succeed at stopping orangutan loss.

### Implications for biodiversity conservation

We showcased how information on deforestation risk and wildlife density can be combined to draw insights into conservation threats and vulnerability assessment. This allows us to understand where areas of high orangutan density might be affected by forest loss and where a reduction in deforestation risk would lead to largest increases in species protection. Such information could be used to direct orangutan conservation efforts, for example by contributing to Population and Habitat Viability Assessments (Utami-Atmoko et al., 2019), national orangutan conservation action plans (Ministry of Environment and Forestry., 2019) or influencing funding or specific interventions across the species range. In future, scenario analysis could help improve landscape-scale planning with largest benefits for orangutan conservation.

The plight of orangutans attracts considerable public interest. The flagship species and their habitat are a focus of research and conservation efforts (Marshall et al., 2016), attracting conservation funding that can help to also protect the habitat of sympatric species. As a consequence of this public attention, however, the available data that underpins the range-wide density distribution model is much better than that available for many other species. In the future, methods that facilitate abundance estimates over large spatial scales, such as integrated modelling that can harness a wider range of data (Bowler et al., 2019) could make abundance estimates more readily available for more elusive or less-well studied species. However, valuable information can also be gleaned from inspecting deforestation risk within species ranges or in combination with occurrence probabilities (e.g., Boitani et al., 2011). This would enable the assessment of more general effects of future forest loss on tropical fauna.

Our findings present a window of opportunity to curb deforestation and its impacts on biodiversity, while highlighting the consequences if we fail to do so. In the context of extensive and rapid changes of land-use, land-cover and climate this century, increasing efforts to further such approaches and to translate them into effective conservation actions is urgently needed to halt wildlife decline in biodiversity hotspots such as Borneo. Ideally, conservation actions now should not only attempt to act on today’s information about deforestation patterns, but also be adaptive to potential changes in drivers and threats.

## Acknowledgements

M.V. thanks Sergio Maroccoli and Ana D. de Lima Voigt for comments on early versions of this manuscript. M.V. and H.K thank the Max Planck Society and Robert Bosch Foundation for funding and support.

## Author Contributions

Conceptualization, M.V., H.M.P., M.A., E.M., J.S., M.J.S., S.A.W., H.S.K., and I.MD.R.; Methodology, M.V., H.M.P., F.W., and I.MD.R.; Software, M.V., F.W., and I.MD.R.; Validation, M.V., I.MD.R.; Formal analysis, M.V.; Resources, D.G., T.S.; Data curation, M.V.; Writing – Original draft, M.V.; Writing - Review & Editing, M.V., H.M.P., M.A., D.G., E.M., T.S., J.S., M.J.S.,S.A.W., F.W., H.S.K., and I.MD.R.; Supervision, H.M.P, H.S.K., and I.MD.R..

## Supporting Information

- S1 Spatial layers of deforestation drivers and processing.
- S2 Deforestation model and calibration.
- Table S1: Overview over best models and predictor effect sizes for each province.
- Table S2: Province area, forest area and projected proportion of forest loss.
- Table S3: Difference between observed and projected annual deforestation rates for the calibration period 2013-2017.
- Figure S1: Proportion of match between observed and cumulative forest loss within the neighborhood of a pixel for Borneo and each province.
- Figure S2: Observed deforestation and projected probability of forest loss across Borneo (2001–2032).
- Figure S3: Summed probability of forest loss and orangutan density across Borneo.
- Figure S4: Map of density of orangutans and map of summed probability of forest loss in provinces.
- Figure S5: Bivariate map of density distribution of orangutans and summed probability of projected forest loss until 2032 in unprotected areas outside of concessions, Lesan-Wehea Landscape and an area in the periphery of Sabangau National Park.

