## Supporting Information for "Deforestation projections imply range-wide population decline for critically endangered Bornean orangutan"

### S1 Spatial layers of deforestation drivers and processing

In the deforestation model, forest loss was parameterized by using a forest cover layer from Gaveau et al., (2018; 2019) incorporating changes based on global forest loss estimates by Hansen et al., (2013) (Figure 2c in main text).

To account for the varying probability of deforestation between areas designated for different land-use types, we included a layer of land-use and management as a predictor of forest loss (Figure 2b in main text, (Santika et al. 2015)). This layer includes forests within protected areas and logging concessions and unconverted forests within industrial timber plantation concessions, industrial oil palm plantation concessions, and forests outside of protected areas and concessions.

We classified the level of protection according to the World Database on Protected Areas (WDPA) (IUCN & UNEP-WCMC 2017). We only considered areas from the WDPA present in the layer by Santika et al. (2015), as these were derived from national data, assumed to be more representative of the situation on the ground. All areas included in both sources and ranked as category 1-3 in the WDPA were combined in one class ('strict conservation'), which represents the highest protection and areas with little to no active human intervention (Dudley 2013). Classes 4-6, where sustainable use can be practiced (Dudley 2013) and areas included as 'not applicable' and 'not reported' (but still included in national land-use planning as protected area) were categorized into a 'sustainable use' class. All areas that were included in Santika et al. (2015), but missing in the WDPA database were classified as 'national' protected areas. They constitute, for example, protection forest (*Hutan lindung*) and wildlife and 'nature reserves' (*cagar alam)* in Indonesia; 'protection forest reserves' and 'wildlife reserves' in Sabah, and protected forests in Sarawak (Santika et al. 2015).

All predictors where clipped with the forest cover in 2000, since the model does not calculate probability of forest loss for pixels deforested before. All layers were converted to the Asia South Albers Equal Area Conic projection and resampled to the same extent and origin at 1 $km^2$ cell size, the highest resolution available for all layers, using bilinear for continuous and nearest neighbor resampling for categorical predictors.

Severe selective logging is detectable as forest loss at a 30 m-resolution. When increasing the spatial grain, the contextual information of local deforestation events is degraded and can be misinterpreted as clear cut deforestation, even when the aggregated area of forest loss is maintained (but see: Amoroso et al. 2018). Thus, when using the terms ‘forest loss’ or deforestation, we acknowledge that we cannot differentiate severe forest degradation over a larger area, for example through intensive logging, from clear-cutting of forest.

Forest loss on Borneo was analyzed within geopolitical units. Province (for Indonesia), state (for Malaysia) and country borders (for Brunei) were downloaded from the Global Administrative Areas database (Global Administrative Areas 2012) and combined within the extent of the island. Analysis excluded Brunei, as important predictors were missing for the country and it does not harbor orangutans.

All spatial manipulations were performed in Python (Python 2016), using gdal (GDAL/OGR Contributors 2017) and numpy (Oliphant 2016) packages, and aggregated, analyzed and visualized in Python, R (R Core Team 2017) and ArcGIS (Esri Inc. 2014).

### S2 Deforestation model and calibration

The model of forest loss for each province and state was adapted from Rosa et al. (2013) and is based on $P_{trloss,x,t}$, the probability that trees in a cell $x$ are lost in a time interval $t$. The probability of loss is defined as a logistic function:

$$Ptrloss_{x,t} = \frac{1}{1 + exp^{-k_{x,t}}} \qquad (1)$$

in which $k_{x,t}$ can range from minus to plus infinity and $P_{trloss,x,t}$ from 0 to 1. We then used linear models to describe $k_{x,t}$ as a function of the predictor variables that affect forest loss at location $x$ and time $t$.

Using a forward stepwise regression, a total of 31 models were fitted to the observed forest loss data (2013 – 2017). Each model differed in the combination of predictor variables that define $k_{x,t}$. The models were fitted using ‘Filzbach’, a freely available library (https://github.com/predictionmachines/Filzbach), which uses a Markov Chain Monte Carlo (MCMC) sampling method to return a posterior probability distribution for each parameter. From this distribution, given a specific parameter combination Θ, the posterior mean and credible interval was extracted. To estimate the parameters, the log-likelihood, a measure of the goodness of fit between the observations and the model predictions, is defined for a particular combination of variables:

$$L(X \vee s, \theta) = \sum log\left(Z_{x,t}Ptrloss_{x,t} + (1 - Z_{x,t})(1 - Ptrloss_{x,t})\right) \qquad (2)$$

in which $Z_{x,t}$ is the observed forest loss at location $x$ and time $t$, and $s$ one of the 31 models considered.

To assess the predictive power gained by adding variables to the model, a cross-validation technique was used. This technique allowed to check how accurately the model predictions compared to a randomly selected subset of 50% of the data that was not used to train the model. This cross-validation is necessary to find models that only comprise predictors with evident predictive ability. After successively adding the variable that resulted in the highest likelihood model, the overall best model (i.e. the one with the maximum test likelihood) was selected from the whole set of models for each province.

The simulations were based on recalculating equation (1) for each time-step, while using a slightly different set of parameter values at each iteration, thereby incorporating parameter uncertainty. These values were drawn from a Gaussian distribution resulting from the MCMC fitting, using the estimated mean and standard deviation for each parameter. As a result we received an updated $P_{trloss,x,t}$ for each individual cell (*x*) in each individual time period (*t*). We subsequently evaluated whether or not the respective pixel was lost, by drawing a random number from a uniform distribution between 0 and 1. We then classified the pixel as lost, if the number was less than the probability of deforestation $P_{trloss,x,t}$. This procedure was repeated for all four time-steps and run multiple times (n = 100 iterations) to assess the uncertainty in model predictions over time. The different iterations were aggregated into the summed probability of deforestation and represent the fraction of simulation runs in which the forest in a pixel in location *x* was lost.

Initial models suggested that the inclusion of a predictor representing the type of soil (mineral or peat), did not significantly improve model predictions. Hence soil types were not included. All predictor variables, except for forest loss, were static, i.e., only one time-step was considered, while forest loss in the neighborhood of a cell was dynamically updated by the model in each time-step.

**Table S1:** Overview over best models and predictor effect sizes for each province. Models were ranked according to their test likelihood and the model with the maximum test likelihood per province is show.

| Province | Sabah | Sarawak | West Kalimantan | South Kalimantan | Central Kalimantan | East Kalimantan | North Kalimantan |
|---|---|---|---|---|---|---|---|
| Test Likelihood[b] | -3,003.5606 | -7,252.6683 | -5,678.1869 | -777.9408 | -7,806.2213 | -4,022.0491 | -2,494.173 |
| Intercept | -2.5678 | -2.5542 | -2.0578 | -2.1246 | -1.644 | -2.2285 | -2.3206 |
| Previous deforestation | 4.2044 | 3.2768 | 3.5371 | 2.6688 | 3.8692 | 4.4006 | 3.4051 |
| Distance to road | -0.0002 | -0.0002 | 0 | 0 | 0 | -0.0002 | -0.0001 |
| Distance to river | 0 | 0 | 0 | 0 | 0 | 0 | - |
| Fire incidence | 0 | 0 | 0 | - | 0.0001 | 0 | - |
| Elevation | - | -0.0007 | -0.0062 | -0.0031 | -0.0097 | -0.0015 | -0.0033 |
| Population density | 0.0001 | - | 0.0002 | - | - | 0 | 0.0002 |
| Strict Protected Area | -1.9465 | -1.6718 | -1.5418 | -1.6142 | -0.2912 | 1.2358 | -1.1241 |
| Sustainable use Protected Area | -1.1166 | -0.5832 | -0.0575 | -0.0001 | -0.2958 | -0.1787 | -0.0488 |
| National Protected Area | -0.7207 | -0.6811 | -0.2723 | -0.2916 | 0.0541 | -0.6829 | -0.5877 |
| Logging Concession | -0.2545 | -0.0838 | -0.2339 | 0.1031 | -0.1263 | -0.2025 | -0.1585 |
| Timber Plantation Concession | -0.0757 | -0.0225 | 0.0083 | 0.0124 | -0.1796 | 0.0916 | 0.1565 |
| Oil palm Plantation Concession | -0.1256 | -0.0346 | 0.0953 | -0.0358 | 0.0338 | 0.0753 | 0.0554 |

**Table S2:** Province area, forest area and projected proportion of forest loss.

| | | **Forest in 2000** | | **Forest in 2017** | | **Forest area in 2032 in km$^2$** | | | **Forest loss 2018 to 2032 in %** | | |
|---|---|---|---|---|---|---|---|---|---|---|---|
| **province** | **Area in km$^2$** | **Area in km$^2$** | **%** | **Area in km$^2$** | **%** | **Median** | **Lower Confidence Interval** | **Upper Confidence Interval** | **Median** | **Lower Confidence Interval** | **Upper Confidence Interval** |
| Sabah | 73,541 | 43,495 | 59 | 37,605 | 51 | 30,747 | 30,571 | 30,861 | 18 | 18 | 19 |
| Sarawak | 123,797 | 78,996 | 64 | 61,900 | 50 | 46,912 | 46,666 | 47,153 | 24 | 24 | 25 |
| West Kalimantan | 146,981 | 69,927 | 48 | 58,841 | 40 | 44,883 | 44,680 | 45,114 | 24 | 23 | 24 |
| South Kalimantan | 36,620 | 8,841 | 24 | 7,556 | 21 | 5,901 | 5,815 | 5,969 | 22 | 21 | 23 |
| Central Kalimantan | 153,568 | 90,471 | 59 | 75,833 | 49 | 53,969 | 53,654 | 54,192 | 29 | 29 | 29 |
| East Kalimantan | 126,783 | 64,781 | 51 | 59,207 | 47 | 49,390 | 49,179 | 49,629 | 17 | 16 | 17 |
| North Kalimantan | 69,840 | 63,145 | 90 | 58,774 | 84 | 53,164 | 53,046 | 53,340 | 10 | 9 | 10 |

**Table S3:** Difference between observed and projected annual deforestation rates for the calibration period 2013-2017 (in comparison to 2000). Annual deforestation rates were averaged over five years, with the exception of maximum observed rate, which is for one year.

| province | Annual deforestation rate | | | | | Difference between observed and projected | | |
|---|---|---|---|---|---|---|---|---|
| | Observed | Maximum Observed | Median projected | Lower Confidence Interval | Upper Confidence Interval | Median | Lower Confidence Interval | Upper Confidence Interval |
| Sabah | 0.76 | 1.01 | 0.84 | 0.8 | 0.88 | 0.08 | 0.04 | 0.12 |
| Sarawak | 1.12 | 1.6 | 1.22 | 1.18 | 1.25 | 0.1 | 0.06 | 0.14 |
| West Kalimantan | 1.18 | 1.58 | 1.51 | 1.47 | 1.55 | 0.33 | 0.29 | 0.37 |
| South Kalimantan | 1.07 | 1.59 | 1.14 | 1.04 | 1.25 | 0.07 | -0.03 | 0.18 |
| Central Kalimantan | 1.21 | 3.3 | 1.51 | 1.47 | 1.56 | 0.3 | 0.26 | 0.35 |
| East Kalimantan | 0.74 | 0.95 | 0.88 | 0.84 | 0.92 | 0.14 | 0.1 | 0.17 |
| North Kalimantan | 0.48 | 0.68 | 0.55 | 0.53 | 0.58 | 0.07 | 0.05 | 0.1 |

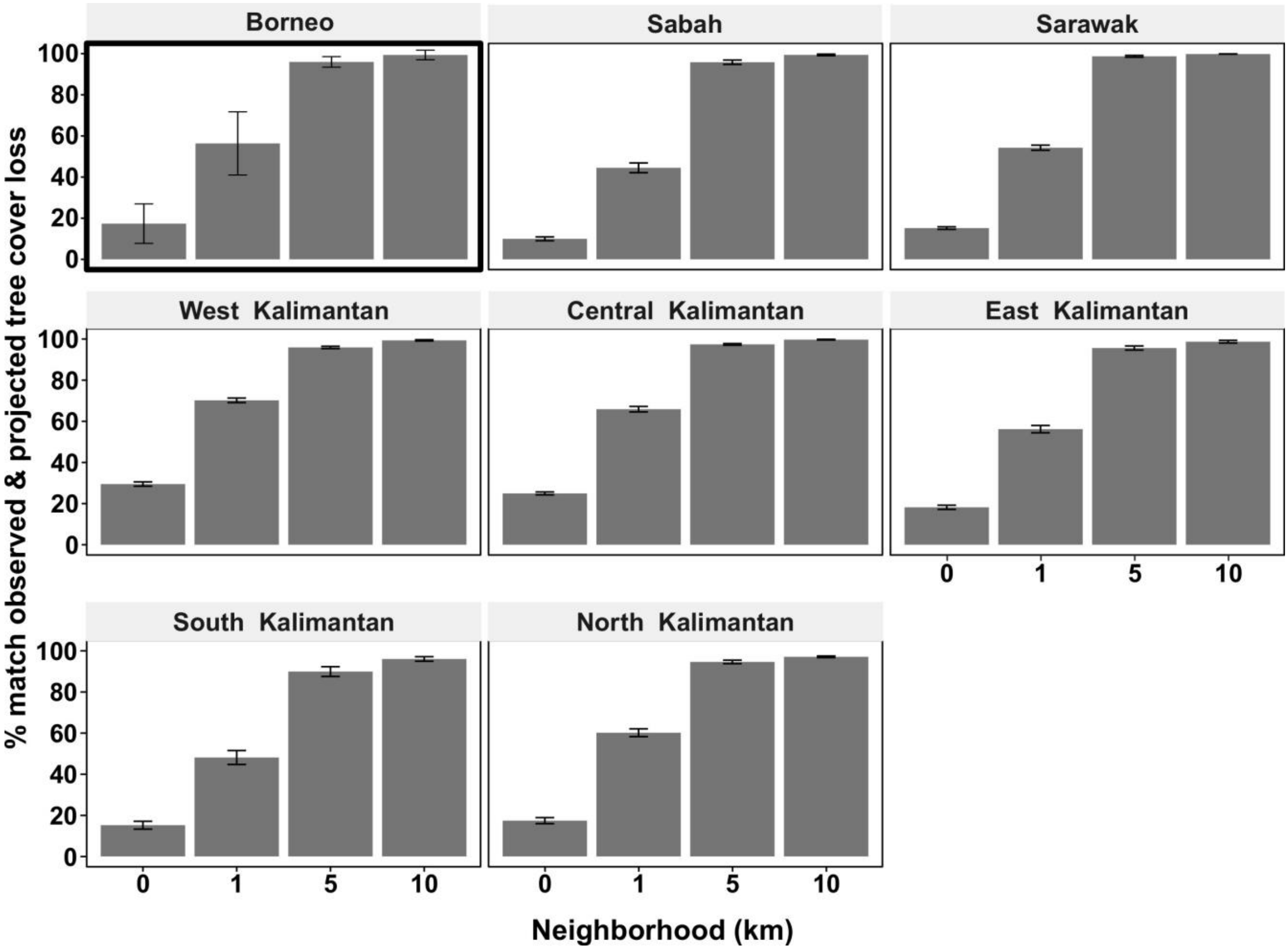


**Figure S1:** Proportion of match between observed and cumulative forest loss within the neighborhood of a pixel for Borneo (pooled over provinces) and each province. Bars show the median across simulations (n=100). Thick outer border indicates the summary plot. The errorbars indicate interquartile ranges.

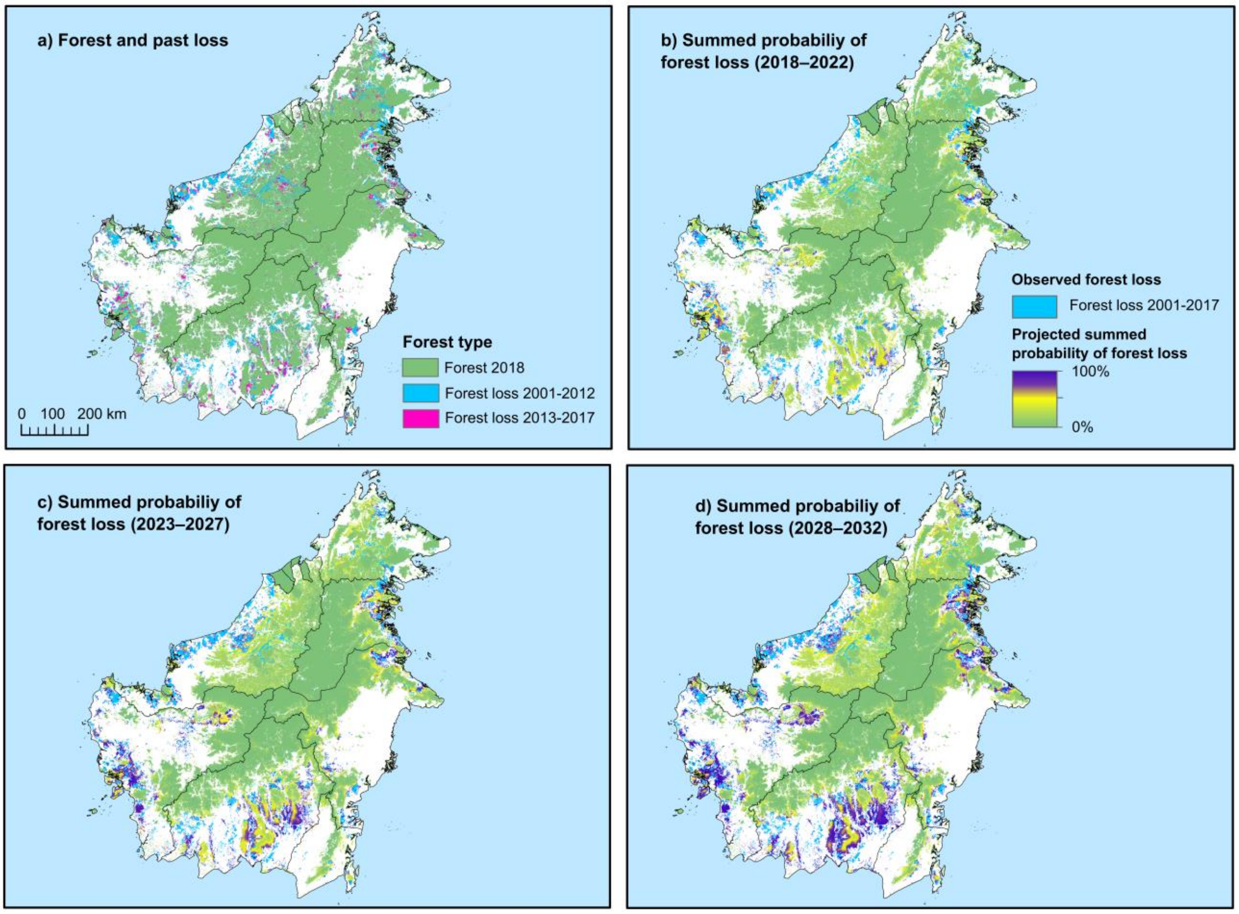


**Figure S2:** Observed deforestation and projected probability of forest loss across Borneo (2001–2032). a) Remaining forest in 2018, past forest loss (2001-2012) and loss in calibration period (2013-2017). b-d) Summed probability of projected forest loss in five-year time steps from 2018 to 2032.

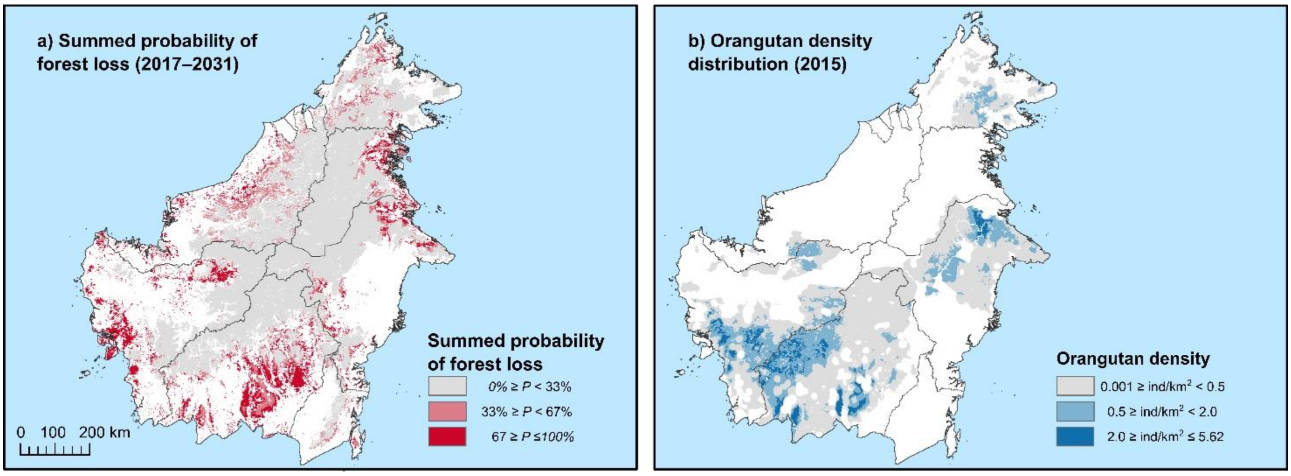


**Figure S3:** Summed probability of forest loss and orangutan density across Borneo. a) The distribution of projected probability of forest loss in three classes for all pixels forested in 2000. b)

Orangutan density distribution in three classes for all pixels with a density higher than 0.001 ind/km$^2$.

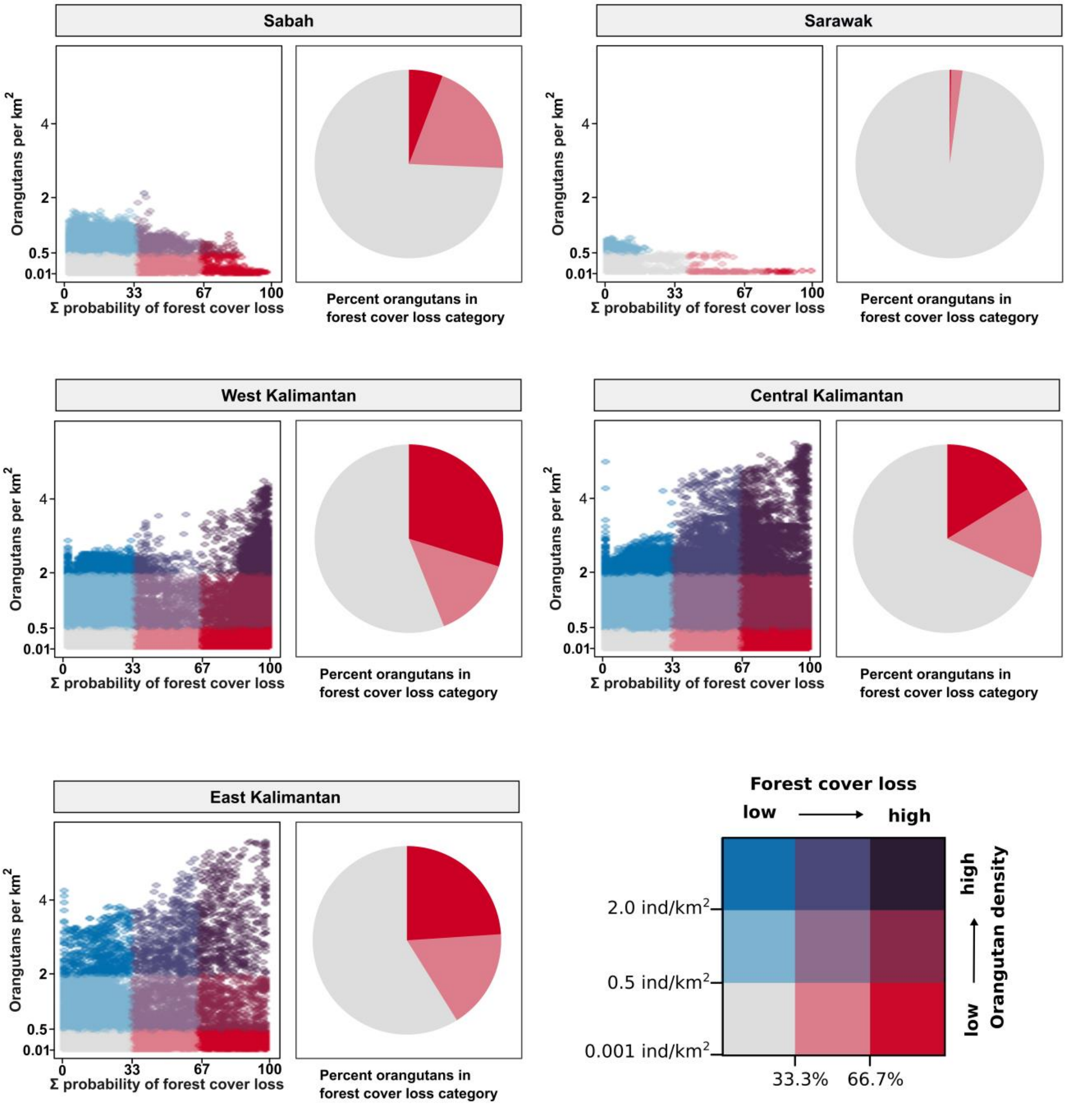


**Figure S4:** Density of orangutans and summed probability of forest loss in provinces. Density of orangutans (blue) and summed probability of forest loss (red). Blue and red shades indicate either factor, intensity corresponding to values, purple hues represent a mix of elevated levels (in maps and scatterplot). The distribution of pixels with respect to the orangutan density per square-kilometer and the summed (Σ) probability of forest loss in scatterplot. The proportion of orangutans in areas with low, medium or high levels of forest loss in pie charts, red shades only. North and South Kalimantan are not shown, as low number of orangutans (<100 individuals) occurred there.

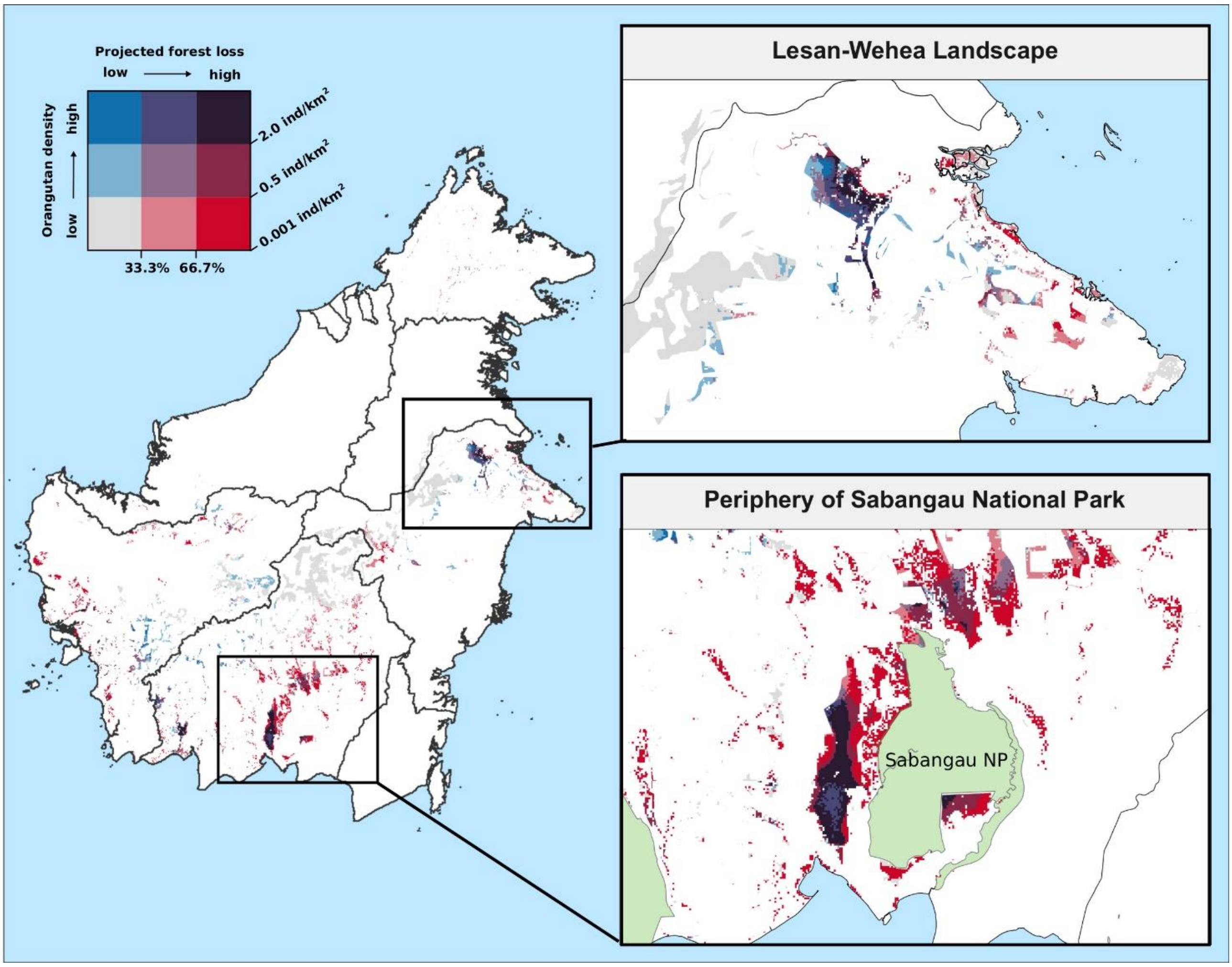


**Figure S5:** Density distribution of orangutans and summed probability of projected forest loss in unprotected areas outside of concessions until 2032. Two areas with especially high orangutan densities at high risk are highlighted in insert maps: Lesan-Wehea Landscape and an area in the periphery of Sabangau National Park (NP), especially in the west of the park.
